# XAMP: debiased dual-engine AI framework enables global discovery of antimicrobial peptides from deep-sea microbiomes

**DOI:** 10.1101/2025.11.20.689422

**Authors:** Bairun Chen, Xinyi Mou, Zhuoxuan Song, Huaying Lin, Tianyi Han, Runze Wang, Hong-Yu Ou, Yu Zhang, Jing Li

**Author notes:** Corresponding author: Jing Li, Yu Zhang. These authors contributed equally: Bairun Chen, Xinyi Mou, Zhuoxuan Song, Huaying Lin.

## Abstract

The global crisis of multidrug-resistant pathogens necessitates innovative antimicrobial peptide (AMP) discovery. However, current AMP predictors are severely compromised by three overlooked systematic data biases, including sequence length imbalance, N-terminal methionine artifacts, and lack of microbial optimization. To overcome these fundamental bottlenecks, we developed XAMP, a dual-engine predictor integrating XAMP-E (ESM-2-based for high accuracy) and XAMP-T (a one-layer Transformer for high speed), trained on rigorously debiased datasets. Consequently, XAMP achieved a median AUC of 0.972, representing an improvement of up to 21.9% in AUC while operating 5 to 40 times faster. Beyond accurate classification, we found that AMPs inherently possess a significantly higher net charge (+3.9 vs. +0.5) despite similar hydrophobicity, along with distinct cooperative patterns such as W-K/R pairs. Applying this pipeline to 238 deep-sea metagenomes (>1000m depth), we identified 2,355 promising AMP candidates from microbial dark matter, and metaproteomic analysis confirmed in-situ expression for a subset of these candidates. Notably, these deep-sea AMPs exhibited a significant 1.53-to 2.93-fold enrichment in K/R/W/Y residues compared to known AMPs. Experimental validation demonstrated that six synthesized peptides exhibit potent, broad-spectrum activity against *ESKAPE* pathogens, with particular efficacy against Gram-negative bacteria that dominate deep-sea ecosystems and pose major clinical challenges. This study establishes that correcting algorithmic data biases is a prerequisite for reliable mining, providing a robust framework to unlock the therapeutic potential hidden within microbial dark matter, offering new avenues to combat the antibiotic resistance crisis.

## Introduction

Antibiotics have been foundational to modern medicine^[1]^, but their overuse and the horizontal transmission of resistance genes have fueled the global crisis of multi-drug resistant (MDR) bacteria^[2]^. By 2050, MDR infections are projected to cause over 10 million annual deaths, surpassing cancer as a leading cause of mortality^[3]^. This crisis demands alternative therapeutics^[2]^, with antimicrobial peptides (AMPs) emerging as promising candidates due to their broad-spectrum activity and unique mechanism of action—targeting bacterial membranes rather than specific genetic pathways, thus minimizing resistance development^[4, 5]^.

Recent breakthroughs in metagenomic analysis and artificial intelligence (AI) have accelerated AMP discovery. Marcel, a random forest-based AMP prediction tool, integrates six local and sixteen global features to enable end-to-end prediction directly from (meta)genomic raw data^[6]^. The deep learning framework c_AMPs-prediction, employs an architecture combining Long Short-Term Memory (LSTM), attention mechanisms, and Bidirectional Encoder Representations from Transformers (BERT) for identifying AMPs within human gut microbiota^[7]^. iAMPCN, a Convolutional Neural Network (CNN)-based model, achieves accurate identification of 22 functional activities by integrating four distinct types of sequence features^[8]^. APG leverages the ESM-2 protein language model as a discriminator to differentiate between real and fake peptides generated by its generator^[9]^.

However, despite the success of tools like Macrel and c_AMPs-prediction, we identify three systematic data biases that severely impair their reliability for mining novel microbiomes:(i). Length distribution imbalance: Most predictors use UniProt-derived negative samples, which lack short proteins, leading to spurious negative correlations between prediction probability and length. Also, APG’s short length focus on 11-30 amino acids (aa) underscore the need for broadly applicable predictors. (ii). N-terminal methionine (Met) artifacts: Mature proteins often undergo N-Met excision, but 92.7% of non-AMPs in c_AMPs’s trainset retain N-Met, distorting feature spaces. (iii). Microbial origin specificity: Existing models lack optimization for microbial AMPs, which dominate deep-sea and extremophilic environments but differ in physicochemical properties (e.g., charge and hydrophobicity) from non-microbial AMPs.

While these methodological challenges persist, AI-driven mining of metagenomic data has yielded substantial breakthroughs in discovering AMPs from diverse biological sources. From the human gut microbiome, Ma et al. identified 2,349 candidate AMPs using an integrated NLP pipeline, with experimental validation confirming activity in 181 of 216 synthesized peptides, showing efficacy even against drug-resistant lung infections in mice^[7]^. Similarly, Xu et al. uncovered a large reservoir of AMPs from engineered environments such as sludge, identifying 27 candidates of which 21 exhibited antibacterial activity^[10]^. On a global scale, Santos et al. established the AMPSphere resource by screening over 150,000 metagenomes and genomes, predicting 863,498 non-redundant AMPs and validating 79 out of 100 tested peptides against clinically relevant pathogens^[11]^. Beyond contemporary ecosystems, pioneering studies have also explored unconventional sources: Guan et al. mined animal venoms, identifying 53 active peptides out of 58 tested^[12]^, while Torres et al. uncovered archaea sins from archaeal proteomes, with 93% of synthesized candidates showing potent antibacterial activity in vitro and in vivo^[13]^.

Despite these advances, one major biome remains largely untapped: the deep-sea microbiome. Characterized by extreme conditions and unique evolutionary pressures, deep-sea microorganisms represent an exceptional resource for discovering AMPs with novel mechanisms and structures. However, deep-sea-derived AMP resources have been significantly under-explored, partly due to technical challenges in sampling, sequencing, and computational prediction (including above three systematic data biases).

To bridge this gap, we present XAMP, a dedicated debiased dual-engine AI framework designed to unlock the therapeutic potential hidden within the “microbial dark matter” of extreme environments (Figure 1). Our approach first addresses fundamental algorithmic artifacts—specifically length imbalance and N-terminal methionine (N-Met) artifacts—through a rigorous data curation pipeline to establish a debiased dataset. Leveraging this foundation, XAMP integrates two complementary engines: XAMP-E (ESM-2-based for high-accuracy semantic capturing) and XAMP-T (Transformer-based for ultra-fast screening), enabling a “consensus screening” strategy that balances precision and efficiency. By deploying this pipeline to 238 deep-sea metagenomes, we constructed a comprehensive Deep-sea AMPs Database containing 2,355 high-confidence candidates. The biological reality of these predictions was substantiated through metaproteomic evidence and experimental validation against *ESKAPE* pathogens, providing a scalable paradigm for AI-driven antimicrobial discovery from unexplored biomes.

**Figure 1.**
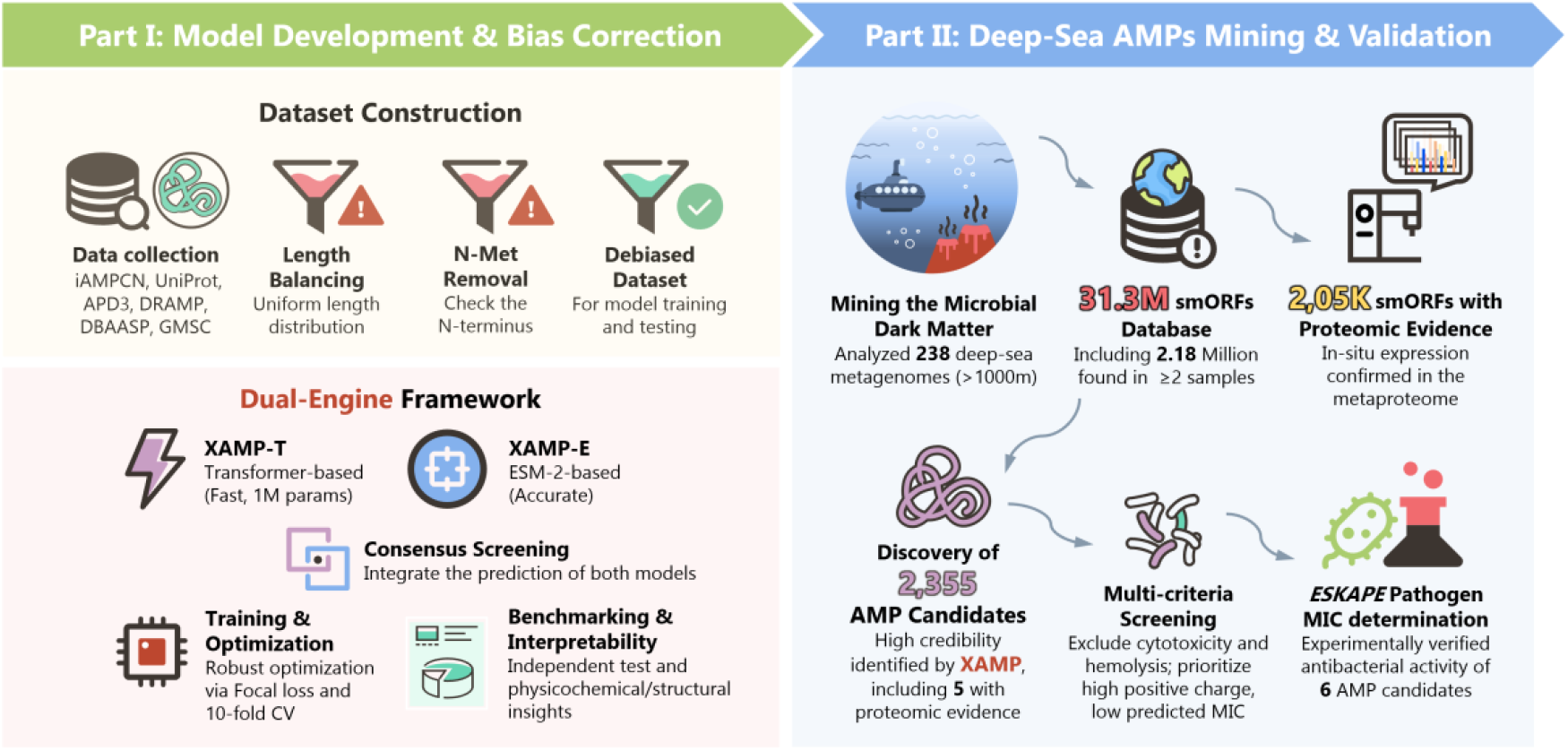
XAMP: A Debiased Dual-Engine Framework for the Discovery of Deep-Sea AMPs. The workflow is divided into two synergistic modules: Part I: Model Development & Bias Correction (left) and Part II: Deep-Sea AMPs Mining & Validation (right). (Part I) We curated a diverse trainset from several public databases. Systematic biases were eliminated via length balancing and N-Met removal to generate a debiased dataset. The dual-engine architecture integrates XAMP-T (a lightweight 1M-parameter Transformer for high-speed screening) and XAMP-E (an ESM-2-based engine for high-accuracy feature representation), optimized through focal loss and 10-fold cross-validation to ensure robustness and interpretability. (Part II) The framework was applied to “microbial dark matter” from 238 deep-sea metagenomes (>1000m depth). From an initial pool of 31.3 million smORFs, XAMP identified 2,355 high-confidence AMP candidates. Candidates were prioritized by excluding cytotoxicity and hemolysis. The pipeline’s reliability was confirmed through metaproteomic spectral evidence (4 in-situ expression matches) and experimental MIC determination, demonstrating potent, broad-spectrum activity of 6 synthesized deep-sea AMPs against ESKAPE pathogens.

## Results

### Addressing Critical Artifacts and False Positive Risk in AMP Trainset

We first conducted an assessment of prevalent prediction tools. Our analysis reveals that the reliability of existing AMP prediction models is substantially compromised by systematic biases present in their trainsets. Specifically, we identified a pronounced length distribution disparity exists between AMPs and non-AMPs in their trainsets (Figure 2A). For example, less than 4% of non-AMPs are shorter than 50 aa in Macrel’s dataset, compared to over 77% of AMPs, leading to a spurious negative correlation between prediction score and peptide length (Figure 2D). Furthermore, the prevalence of N-terminal methionine (N-Met) in non-AMP sequences creates an artifact that inflates model performance illusorily. Since most non-AMPs in trainsets retain this translation-initiation residue (92.7% in c_AMPs; 97.3% in iAMPCN; Figure 2B)—unlike mature proteins in vivo^[14]^—models learn to misuse its presence as a simple marker for negative classification. This results in artificially optimistic performance during validation, which is exposed as a sharp rise in false positive rates when models are tested on debiased data with N-Met removed (Figure 2C, E).

**Figure 2.**
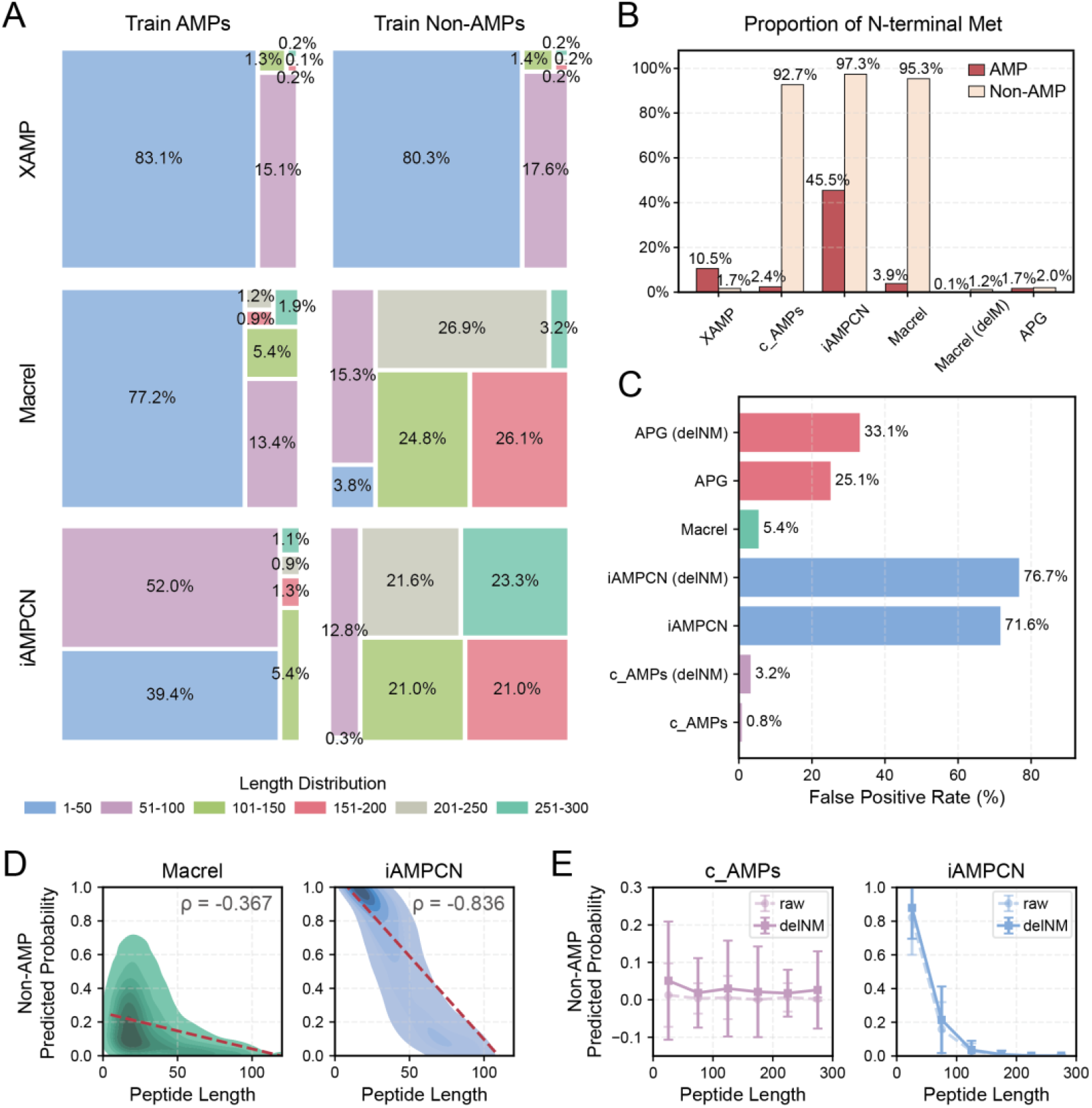
Systematic trainset data biases impair existing AMP prediction reliability. (A). Length distribution disparities between AMPs and non-AMPs in trainsets of major predictors. (B). Proportion of N-Met in AMPs and non-AMPs across trainsets. (C). False positive rate (FPR) comparison on our debiased testset. “delNM” indicates using N-Met-removed non-AMPs as model inputs. (D). Spurious negative correlations between prediction probability and peptide length in Macrel (ρ = -0.367) and iAMPCN (ρ = -0.836) on our debiased testset. (E). Prediction probability of non-AMPs by c_AMPs and iAMPCN before/after N-Met deletion on our debiased testset.

To overcome these limitations, we constructed a rigorously curated benchmark dataset by balancing AMP/non-AMP length distributions and removing N-Met initiation residues from non-AMPs (Figure 2A, B; see *Methods*). The resulting dataset contains more complex sequence characteristics that challenge conventional model architectures, motivating the development of our dual-engine framework, XAMP. By integrating a fine-tuned ESM-2 model for residue-level semantic embedding and a Transformer module for long-range context modeling, XAMP improves the generalization capacity for unbiased AMP recognition.

### Development of XAMP for Mining AMPs in Deep-Sea Microorganisms

We constructed a comprehensive benchmark dataset (“Mix”) totaling 253,203 sequences, which included 246,007 of unannotated origin (13,967 AMPs; 232,040 non-AMPs) and 7,196 of bacterial origin (598 AMPs; 6,598 non-AMPs) (Figure 3A). Analysis of N- and C-terminal motifs revealed distinct compositional patterns: AMPs of unannotated origin were enriched with positively charged lysine (K) and arginine (R) residues, whereas bacterial AMPs were predominantly composed of glycine (G). Non-AMPs were generally characterized by hydrophobic and polar residues (e.g., leucine (L), alanine (A), and serine (S)), consistent with background frequencies in source databases.

**Figure 3.**
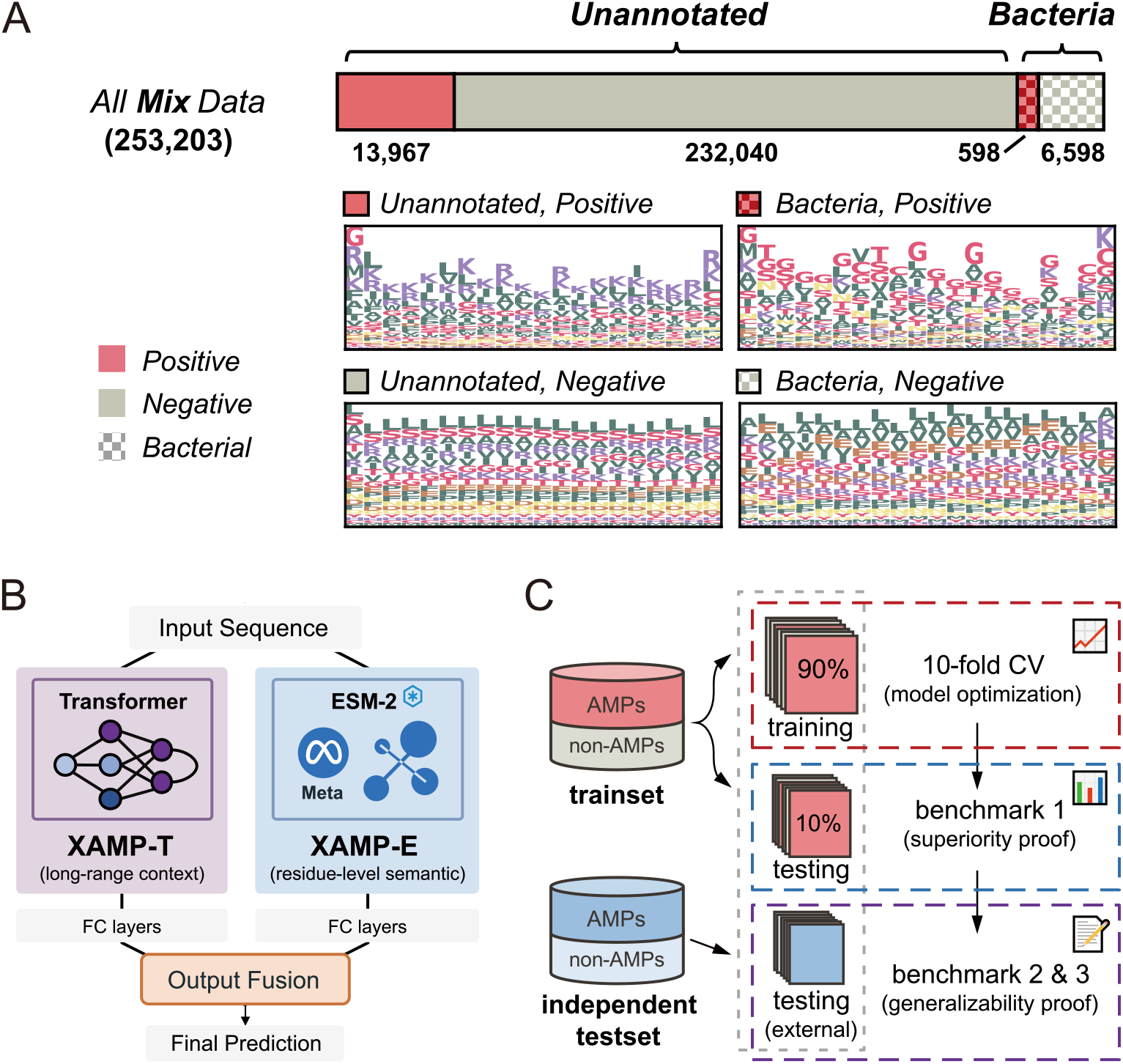
Integrated framework for AMP prediction and analysis. (A). Data composition and motif characterization. Bar plot displays the distribution of sequences in the ‘Mix’ dataset. Sequence motifs are shown for N- and C-terminal regions. (B). Dual-engine architecture of XAMP. The framework integrates two complementary modules: (i) XAMP-T, a Transformer-based encoder for end-to-end sequence classification, and (ii) XAMP-E, which leverages pre-trained ESM-2 embeddings for enhanced feature representation. (C). Data partitioning strategy. The Trainset was split into training and testing sets, with training employing 10-fold cross-validation. Model performance was evaluated on both internal and external independent testsets.

To effectively leverage this data, we developed XAMP, an integrated dual-engine prediction framework (Figure 3B). This architecture combines two complementary approaches: (i) XAMP-T, an end-to-end classifier built on a Transformer encoder followed by fully-connected (FC) layers for deep feature extraction and classification; and (ii) XAMP-E, which augments sequence representation by incorporating embeddings from a pre-trained ESM-2 model, which are then processed through dedicated FC layers. This dual-model strategy provides two application strategies to balance precision and efficiency: using the integrated model that selects the lower prediction probability from both engines to minimize false positives, or employing the faster XAMP-T individually for rapid large-scale screening. This design ensures optimal performance while retaining flexibility for diverse scenarios.

Following deduplication, the dataset was rigorously curated: we balanced AMP/non-AMP length distributions and systematically removed N-Met initiation residues from non-AMPs. Then, sequences were partitioned into trainset (90%) and testset (10%) (Figure 3C). Model hyperparameters were optimized via 10-fold cross-validation, and final performance was rigorously assessed on both the internal test set and an independent external validation set to confirm robustness and generalizability. Detailed dataset statistics and methodological procedures are provided in Supplementary Table 1 and the Methods section, respectively.

### The Dual-engine XAMP Framework Enables Accurate and Efficient AMP Identification

We first assessed the impact of training data through an ablation study. The superior performance of models trained on the “Mix” dataset highlighted the importance of broad data integration for generalization (Supplementary Table 2). Consequently, all subsequent analyses used the ***mix*** models.

In benchmarking, XAMP demonstrated superior predictive accuracy, improving precision by 40.0%-89.1%, AUC by 8.22%-21.9% and AUPRC by 20.2%-68.6% over baseline models on testset (Figure 4A). This advantage held on external, rigorously designed testsets (Figure 4B-C), confirming excellent generalization.

**Figure 4.**
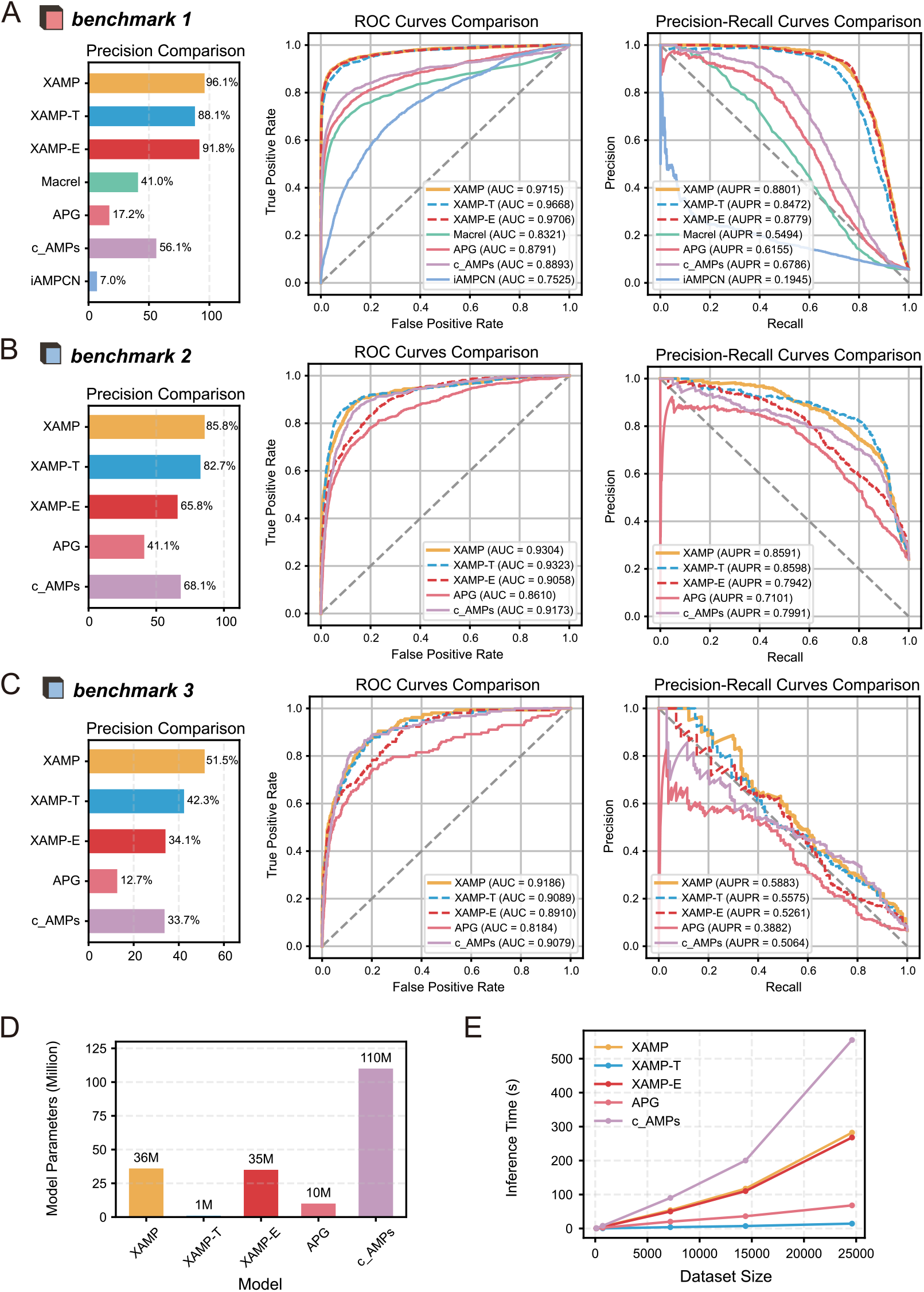
Benchmarking results of the XAMP framework for AMP identification. (A) Predictive performance on the testset (Mix). The precision-recall (PR) curves and receiver operating characteristic (ROC) curves of XAMP and baseline models are compared, with XAMP showing superior predictive accuracy. (B-C) Generalization performance on two independent testsets (Xiao 1 and Xiao 2, respectively). The consistent performance across external datasets confirms the robust generalizability of the XAMP framework. (D). Comparison of model complexity in terms of trainable parameters. The XAMP-T variant demonstrates a highly efficient architecture with only ∼1 million parameters, significantly fewer than other deep learning-based AMP predictors. (E). Inference speed measured on datasets of varying sizes. Together, these results demonstrate the framework’s dual-engine design, which combines high predictive accuracy with exceptional computational efficiency.

Beyond predictive accuracy, a critical advantage of the XAMP framework lies in its flexible and efficient design. XAMP-T required only 1 million parameters (Figure 4D) and operated 5 to 40 times faster than other deep learning models across datasets of varying sizes (Figure 4E). This combination of the high-performance, feature-rich XAMP-E engine and the ultra-efficient XAMP-T engine makes the framework uniquely suited for a wide range of scenarios, from detailed analysis to large-scale screening.

### Model Reveals Physicochemical Basis of AMP Antibacterial Activity

Interpretability analyses show that XAMP’s predictions are grounded in sound physicochemical principles. UMAP visualization confirms that both XAMP-T and XAMP-E effectively separate AMPs from non-AMPs, with separation sharpening after the final layers, indicating refined feature learning (Figure 5A).

**Figure 5.**
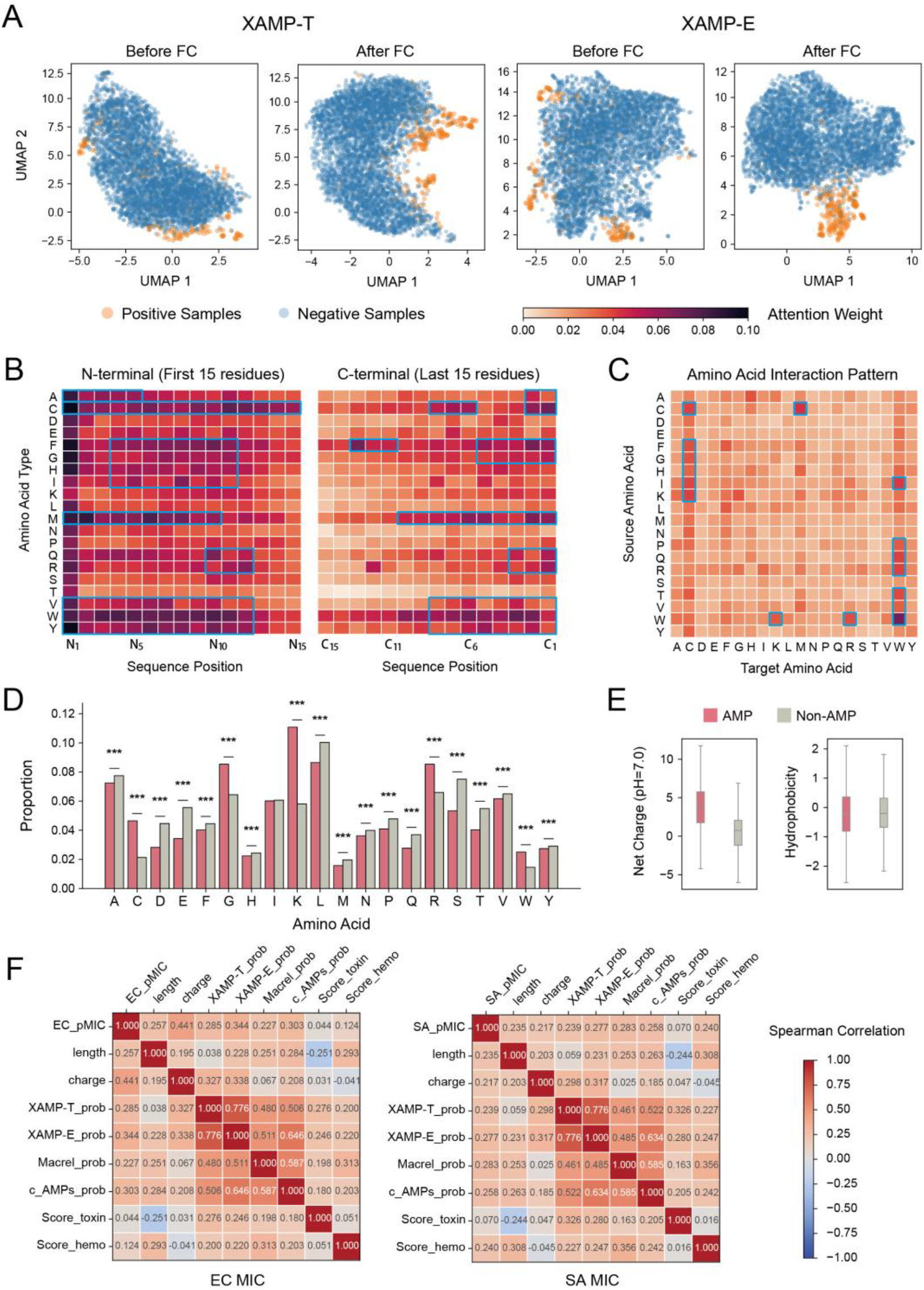
Interpretable features underlying AMP recognition by XAMP. (A)UMAP visualization of the latent space before and after the final network layers. The refined separation of AMPs (orange) from non-AMPs (blue) after the final layers demonstrates effective feature learning. (B)Sequence-level attention map from XAMP-T, identifying critical residues, particularly at the N- and C-terminal (deeper red indicates high attention). (C)Residue interaction matrix revealing cooperative amino acid patterns (e.g., W-K/R pairs), with exemplary strong correlations marked. (D, E) Comparative analysis of amino acid composition (D) and key physicochemical properties (E) between AMPs (red bars) and non-AMPs (gray bars). ‘*’ denote statistically significant differences. Notably, AMPs exhibit a significantly higher net charge despite similar overall hydrophobicity. (F) Spearman correlation matrix between model prediction scores and key peptide properties, including antimicrobial potency (pMIC, defined as -log_10_(MIC)), toxicity, and hemolytic potential. The color gradient represents the correlation coefficient magnitude.

Attention analysis in XAMP-T reveals a focus on N/C-terminal residues and preferences for cationic residues K/R alongside hydrophobic residues A/I/V/F/W/Y. This preference matches conserved physicochemical features of natural AMPs and prior residue enrichment findings^[15, 16]^. Beyond these recognized patterns our model further identifies obvious residue-residue cooperativity represented by combinations such as W-K/R (Figure 5B, C). These newly captured interactive features have rarely been systematically characterized in conventional AMP sequence analyses. The learned patterns overall align with the significant physicochemical differences between AMPs and non-AMPs^[17]^, notably a much higher net charge (+3.9 vs. +0.5) and distinct amino acid enrichment despite similar overall hydrophobicity (Figure 5D, E).

The prediction score of XAMP-E shows a moderate correlation with the MIC for E. coli (Spearman’s r=0.344) from BERT-AmPEP60^[18]^, exceeding other models (Figure 5F). More critically, the moderate correlations between model scores and toxicity/hemolysis metrics (r=0.20-0.33) highlight that potent AMP candidates may carry safety risks, underscoring the necessity of integrating safety screening after activity prediction.

### Mining and Characterization of Deep-Sea AMPs

A total of 31,364,546 non-redundant small proteins comprised of 10 to 100 aa were predicted from 238 deep-sea metagenome samples spanning eight distinct marine habitat types (Figure 6 A, B). To enhance reliability, only sequences detected in more than one sample (2,183,935 sequences in total) were retained and subsequently screened for AMPs. Using the XAMP prediction model, a high-confidence set of 2,355 AMPs was identified, which constitute the Deep-sea AMPs Database (Supplementary Table 3).

**Figure 6.**
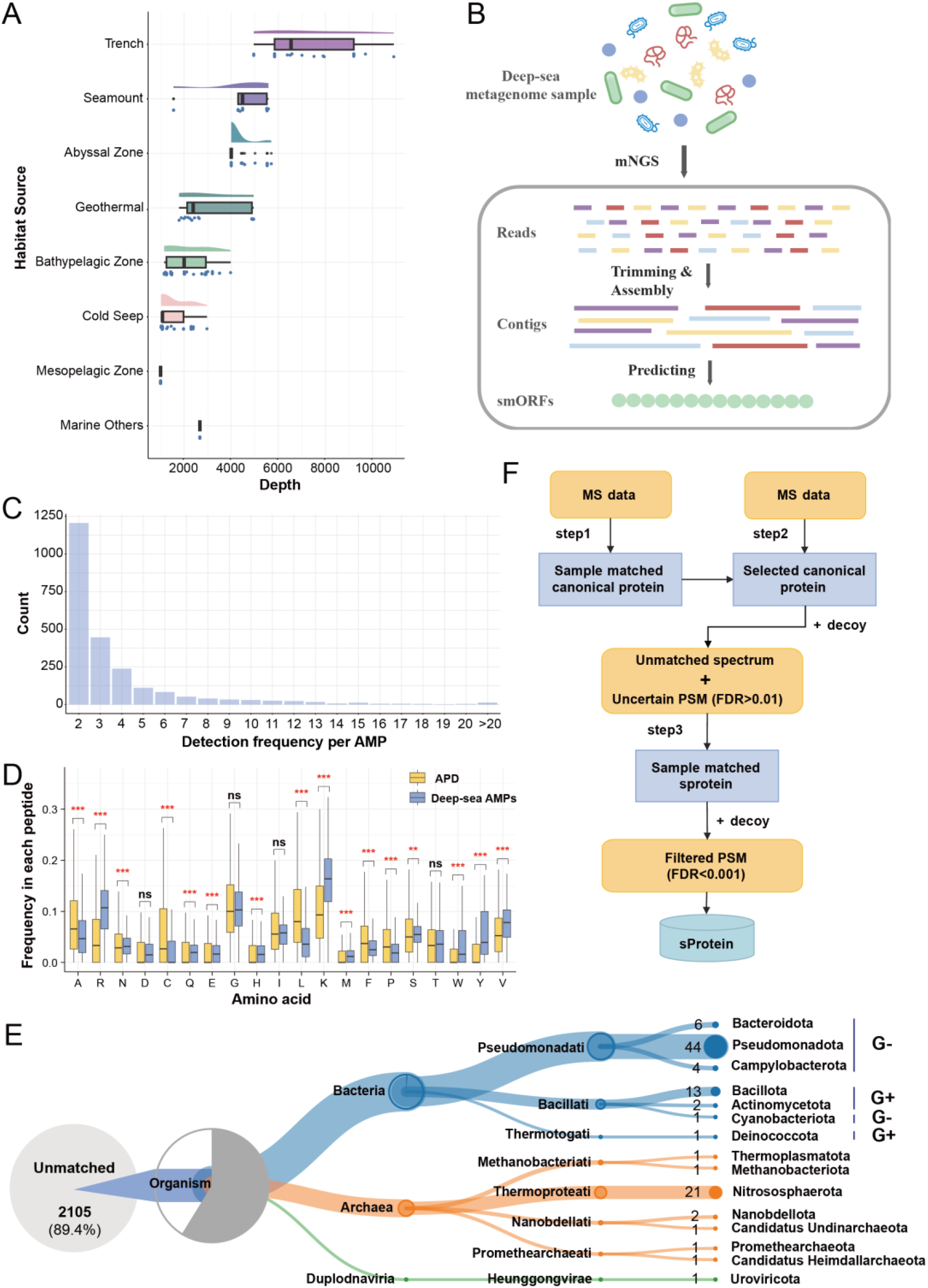
Systematic mining and validation of AMPs from deep-sea microorganisms. (A). Depth Distribution of AMPs across 238 metagenomic samples from diverse marine habitats. (B) Workflow for predicting small open reading frames (smORFs) from metagenomic assemblies. (C). The number of times each antimicrobial peptide from the Deep-sea AMPs Database appeared in the metagenome samples. (D). Comparison of amino acid composition between the Deep-sea AMPs Database and the public APD database. (Wilcoxon rank-sum test with Benjamini-Hochberg correction, ns: not significant, ***:P.adj < 0.001, **:P.adj<0.01, *:P.adj<0.05). (E). Taxonomic lineage of the deep-sea AMPs as determined by Unipept. G+, Gram-positive bacteria; G-, Gram-negative bacteria. (F). Three-step metaproteomic search strategy for validating the expression of predicted small proteins (sProteins) from mass spectrometry (MS) data.

Analysis of the distribution of these AMPs across metagenomic samples revealed distinct ecological patterns: 1,206 were found in exactly two samples, while 12 were detected in over 20 samples, suggesting that a specific subset possesses broad ecological prevalence (Figure 6C). Significant difference was observed in the amino acid composition between the Deep-sea AMPs Database and the public APD database (Wilcoxon rank-sum test, Figure 6D). Specifically, the deep-sea AMPs exhibited a more prominent enrichment (1.53-to 2.93-fold) in K/R/W/Y residues. Consistent with prior findings on deep-sea AMPs^[19, 20]^, this enhanced enrichment likely provides a structural basis for their potent activity against bacteria adapted to high-salt or high-pressure deep-sea environments.

Taxonomic classification via Unipept^[21]^ revealed that 89.4% of the AMPs were unassignable, indicating they originate from unexplored microbial dark matter, likely deriving from novel, unannotated smORFs (Figure 6E). Notably, among the classifiable bacterial-derived AMPs, Gram-negative bacteria were dominant, accounting for 75.0% of this subset. This forms a striking contrast to existing public databases, where microbial AMPs are predominantly derived from Gram-positive bacteria (Supplementary Figure 1). Consequently, our findings highlight the unique value of the deep-sea microbiome as a previously underexplored reservoir, effectively filling a critical gap in the antimicrobial peptide sequence space associated with Gram-negative bacteria.

To assess the in-situ expression of these predicted AMPs, we analyzed deep-sea metaproteomic data using a three-step database search strategy against customized databases built from sample-matched metagenomic ORFs (Figure 6F). From 2,057 identified microbial small proteins, XAMP predicted 5 to possess antimicrobial activity. Crucially, 4 of these overlapped with candidates in our Deep-sea AMPs Database, providing direct proteomic evidence for the in-situ biosynthesis and expression of these computationally predicted smORFs (Supplementary Table 4, Supplementary Figure 2).

### In Vitro Antimicrobial Activities of Deep-Sea AMPs Against *ESKAPE* Pathogens

To validate our computational predictions, we selected seven candidate AMPs from deep-sea metagenome data for chemical synthesis. Six of these were successfully synthesized (Supplementary Table 5), while one failed to synthesize. We evaluated the in vitro efficacy of these six AMPs by determining MICs against five *ESKAPE* pathogens, including four Gram-negative (*E. coli, A. baumannii, P. aeruginosa, K. pneumoniae*) and one Gram-positive (*S. aureus*) strains.

To elucidate the structure-function relationship of the identified peptides, we employed AlphaFold3^[22]^ for 3D structure prediction. The resulting models revealed distinct folding patterns with diverse secondary structure elements (Figure 7A), indicating potential variations in their mechanisms of antimicrobial action.

**Figure 7.**
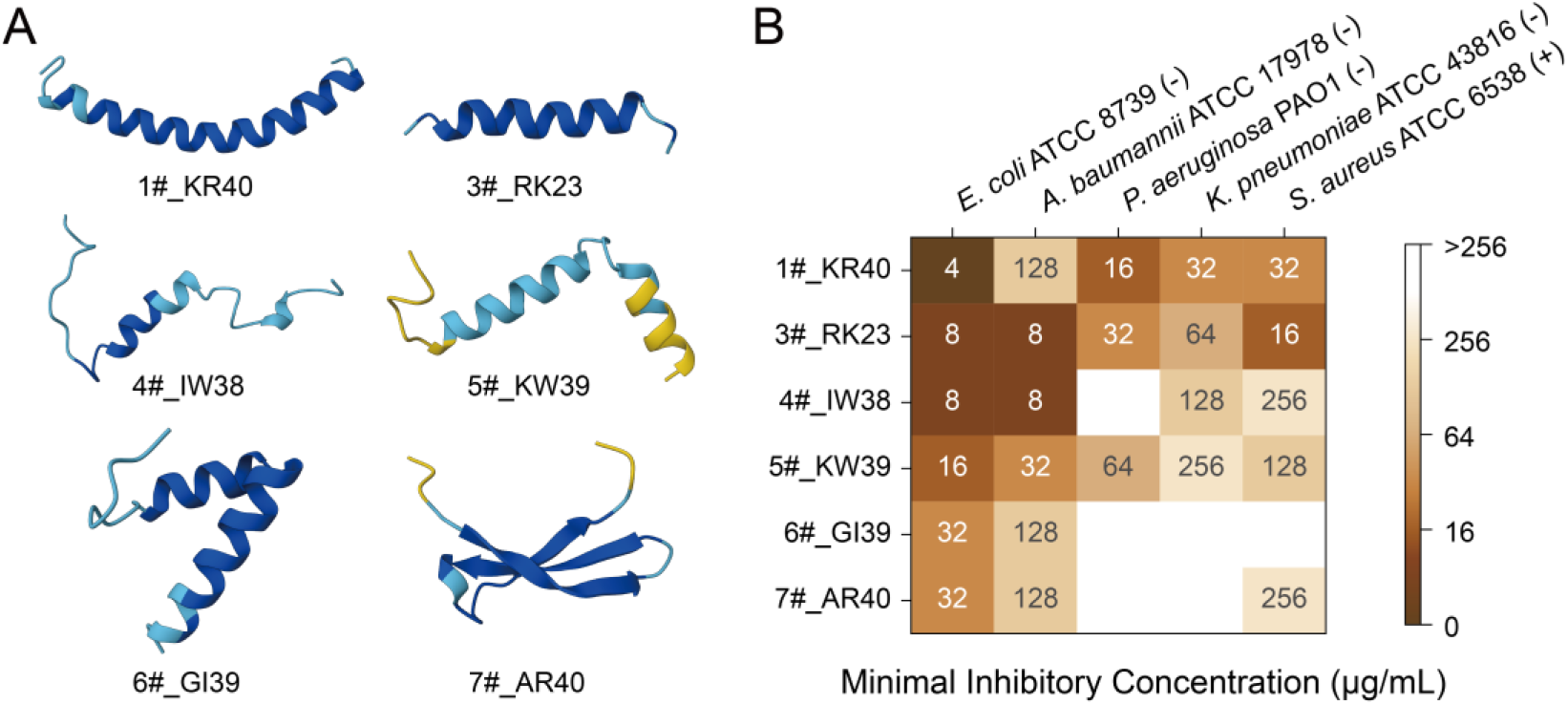
Structural and functional characterization of synthesized deep-sea AMPs. (A). AlphaFold3-predicted 3D structures of six successfully synthesized AMPs. (B) Heatmap of minimum inhibitory concentrations (MICs) against five ESKAPE pathogens. Gram-positive (+) and Gram-negative (−) bacteria are indicated as such (top).

MIC assays (Figure 7B) revealed broad-spectrum antimicrobial activity for all six AMPs, each effective against at least two pathogens. Crucially, the identified AMPs exhibited superior efficacy against Gram-negative bacteria, which are the dominant taxa in deep-sea ecosystems. This result highlights an important “environment-clinic” link: the evolutionary adaptation of deep-sea microbial AMPs to combat Gram-negative competitors translates effectively into clinical potential against challenging nosocomial Gram-negative pathogens. Peptides 3#_RK23 and 4#_IW38, for instance, demonstrated potent activity against *A. baumannii* with an MIC of 8 μg/mL, and also inhibited *K. pneumoniae* ATCC 43816, a strain resistant to multiple antibiotics clinically, validating the pipeline’s ability to discover clinically relevant candidates.

## Discussion

The global antimicrobial resistance crisis necessitates innovative therapeutic strategies. Here, we present XAMP, a dual-engine deep learning framework that overcomes critical data biases in AMP prediction to enable systematic discovery of bioactive peptides from deep-sea microbiomes—an underexplored resource with high therapeutic potential.

We address key methodological bottlenecks in existing AMP prediction. We identified and corrected systematic data biases including sequence length imbalance and residual N-Met artifacts, the primary cause of false positives in metagenomic mining. Built on a debiased dataset, XAMP achieved ∼10% higher AUC than state-of-the-art tools, shifting model learning from dataset-specific noise to authentic biological features. Its dual-engine design balances high-precision experimental validation and ultra-fast large-scale screening, providing a practical tool for microbial research.

This study unlocks the potential of deep-sea microbial dark matter for antibiotic discovery. We identified 2,355 high-confidence AMPs predominantly from unannotated smORFs, and metaproteomic evidence confirmed their in-situ expression, validating the biological credibility of computational smORF predictions.

These deep-sea AMPs exhibit characteristic enrichment of K/R and W/Y, and hold potent activity against drug-resistant Gram-negative *ESKAPE* pathogens. This signature matches the dominance of Gram-negative bacteria in deep-sea ecosystems, indicating extreme environments act as valuable reservoirs for lead peptides targeting refractory Gram-negative infections, aligning with WHO priority needs for anti-resistance therapeutics.

Our study has several limitations warrant consideration. The lower generalizability of bacteria-specific models highlights the challenge of leveraging niche biological data without sufficient sample size or specialized algorithmic design. Additionally, our analysis may underrepresent ultra-short AMPs due to prediction length cutoffs, and current validation is constrained by limited deep-sea proteomic datasets. Furthermore, the in vitro validation was conducted against a limited panel of *ESKAPE* pathogens, more extensive antibacterial assays employing a broader spectrum of bacterial lineages, including diverse clinical isolates, are warranted to substantiate the observed antimicrobial profiles. The study also lacks investigation into the precise mechanisms of action for the active peptides. More critically, all validation was performed in vitro; efficacy and safety within a complex physiological environment remain unevaluated. Finally, while computational screening was used, experimental assessment of cytotoxicity and hemolytic activity for the validated peptides is absent.

Future efforts should focus on expanding taxonomic diversity in training data, developing few-shot learning techniques for specialized prediction tasks, and integrating structural features to enhance prediction accuracy. Scaling metaproteomic validation and conducting systematic functional characterization will further accelerate therapeutic development.

In summary, this study establishes a robust, debiased pipeline for targeted antimicrobial discovery. By combining debiased deep learning with multi-omics validation, our work provides a scalable framework for addressing the urgent threat of drug-resistant pathogens through the rational exploration of microbial dark matter.

## Methods

### Dataset construction

The unannotated-origin training dataset was constructed by collecting antibacterial and antifungal AMPs from iAMPCN’s positive dataset, which was curated from 13 public sources and 25 prior model training sets. Sequences were filtered to ≤ 300 aa, yielding 13,967 peptides^[8]^. For the generation of negative samples at a 20:1 ratio, two steps were employed: (i) Short proteins were extracted from UniRef90, excluding AMP-related annotations such as antimicrobial, antibacterial, antifungal, anticancer, antiviral, antiparasitic, antibiotic, antibiofilm, and effector; (ii) The length distribution of negative samples was matched to that of positive samples. Given the limitations of UniRef, sequences < 10 aa were supplemented from UniProt. After the removal of duplicates, a total of 232,245 negative sequences (≤ 300 aa) were obtained. Notably, for all negative samples in this dataset, the methionine at the N-terminus was removed during the preprocessing stage to ensure consistency in sequence characteristics.

For the bacterial-origin training data, experimentally validated antibacterial/antifungal AMPs were sourced from APD3^[23]^, DRAMP^[24]^, and DBAASP^[25]^, resulting in 598 peptides. The negative samples were composed of: (i) Short non-AMP annotated bacterial proteins from UniProt, with their lengths stratified to match those of positive samples; (ii) Supplemental sequences from the Global Microbial smORFs Catalogue (GMSC)^[26]^ to address the scarcity in UniProt. This process generated 6,598 negative sequences (≤ 300 aa). During the preparation of these negative samples, the N-terminal methionine was also systematically removed. The above two datasets together constitute our mix training dataset.

An independent test set was constructed using Xiao et al.’s benchmark dataset^[27]^. Only antibacterial and antifungal AMPs were retained, and all sequences overlapping with our mix training dataset were removed to avoid evaluation bias. For the negative samples within this test set, the N-terminal methionine was similarly removed to maintain the same preprocessing standard as the training datasets.

### Model development

#### Peptide representation

Initially, the character ‘*’ was padded to the right end of each peptide sequence to achieve a maximum length of 300. Subsequently, we designed tokens for 20 common amino acids, the padding character ‘*’ and the unknown amino acid symbol ‘X’. Ultimately, each token in the peptide sequence was mapped to a distinct integer value, with ‘X’ corresponding to 0, ‘Y’ to 1, ‘S’ to 2, and so forth, up to ‘*’ which was assigned the value 21.

#### XAMP-T architecture

The XAMP-T model is composed of an embedding layer, a positional encoder layer, a single transformer layer, and two FC layers. The processed sequences are padded/truncated to 300 aa using the ‘*’ character, with a masking mechanism applied to distinguish valid sequence content from padding.

The embedding layer converts each token into a 128-dimensional word vector, with its parameters being optimized during the training phase. Sinusoidal positional encoding is employed to encode the positional information of the input sequence. The encoded sequence is subjected to a self-attention layer with eight heads that incorporates the padding mask to exclude ‘*’ symbols from attention calculations.

Subsequently, all word vectors, excluding the padding symbol ‘*’, are averaged to yield a 128-dimensional feature vector that characterizes the peptide sequence information. This 128-dimensional feature vector is then fed into two FC layers, the first with 1024 neurons and the second with 256 neurons. Following these two FC layers, batch normalization is applied to accelerate the convergence of the neural network. Both layers utilize the Rectified Linear Unit (ReLU) as their activation function. In the final stage, the prediction score is computed by the output of the final dense layer, which contains a single neuron. The sigmoid function serves as the activation function, confining the output to the range [0, 1]. A score of 0 indicates that the peptide is classified as non-AMP, whereas a score of 1 signifies an AMP.

### XAMP-E architecture

The XAMP-E model leverages the pre-trained ESM-2 protein language model (version: *esm2_t12_35M_UR50D*)^[28]^. Feature representations extracted from ESM-2 are fed into a stack of FC layers for AMP/non-AMP classification. During training, the weights of the ESM-2 component are frozen, and only the parameters of the FC layers are updated.

#### Loss function

Given the imbalance in the datasets for antimicrobial functional activities, the focal loss function was adopted as follows:

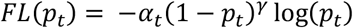

Where α_*t*_ is a hyperparameter that modulates the importance of samples in the positive and negative classes, while *p*_*t*_ represents the probability estimate produced by the model. The term (1 − *p*_*t*_)^*γ*^ acts as a modulating factor that adjusts the impact of easily classified samples. In this implementation, the values of α_*t*_ and *γ* were fixed at 1 and 2, respectively.

#### Training process

The model development adhered to a rigorous training protocol designed to prevent data leakage and ensure robust generalization. The complete dataset was initially stratified into training set (90%) and test set (10%), preserving class distributions in both subsets. Within the training set, a 10-fold cross-validation framework was employed for simultaneous optimization of model architecture, hyperparameters, and training epochs. For each fold, training was strictly capped at 50 epochs, while 90% of the training subset was used for parameter updates while the remaining 10% served as validation monitors, with early stopping automatically triggered if validation AUC plateaued for five consecutive epochs. Using these cross-validated parameters, the model underwent final training on the entire training set. Technical execution employed the AdamW optimizer (weight decay λ=0.01, batch size 128) within PyTorch 2.3.1, with CUDA 12.1 acceleration. This multi-barrier approach effectively decoupled model development from final evaluation, maintaining the integrity of performance metrics.

#### Evaluation metrics

We utilized a diverse set of evaluation metrics to assess the model’s performance, which include the area under the receiver operating characteristic curve (AUC), the area under the precision-recall curve (AUPRC), accuracy, precision, recall, and the false positive rate (FPR):

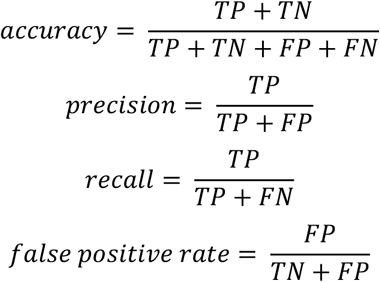

where TP represents the number of true positive instances, FP represents the number of false positive instances, TN represents the number of true negative instances, and FN represents the number of false negative instances.

### Collection of deep-sea metagenome and metaproteome

In this study, we selected and downloaded 238 datasets with sampling depths greater than 1,000 meters through the sample view interface of MASH-Ocean^[29]^ (as of June 2024), which is a platform designed for the systematic and quantitative analysis of microbial communities in marine ecosystems. Given the current challenges in protein sampling and extraction from deep-sea microbial communities, only three deep-sea metaproteomic datasets were retrieved from the PRIDE database for this study. Dataset PXD010074^[30]^ was obtained from hydrothermal vent sites. Dataset PXD009105^[31]^ was collected from vent chimneys. Dataset PXD034421^[32]^ collected 61 metaproteomics samples from 22 stations between 5 m and 4000 m depth from the major ocean basins, and we selected 13 samples with paired metagenomic sequences from > 1000m depth.

### Construction of a deep-sea small protein database

We performed smORF prediction on 238 deep-sea metagenomic sequencing datasets. Raw reads were filtered using Trimmomatic (v0.39), followed by assembly with MEGAHIT (v1.2.9) using the parameter set “--presets meta-sensitive.” Gene prediction was conducted using a modified version of Prodigal with the “-p meta” parameter, targeting smORFs ranging from 33 to 303 bp in length. Finally, redundant small protein sequences were removed to obtain a non-redundant smORF dataset.

### Strategy for deep-sea small protein database search

The construction of the sample-matched deep-sea metaproteomic search databases followed a similar workflow described the previous section, while the upper limit of the ORF length was not set to get both long and small ORFs. In the identification of small proteins from deep-sea metaproteomic data, we implemented a three-step search strategy combined with a target-decoy approach to balance proteome-wide coverage and false positive control.

In the first step, MS-GF+ was used to search against a conventional long protein database (length > 100 aa) without the inclusion of a decoy database. In the second step, the matched long protein entries from the initial search were used to construct a refined target-decoy database, and false discovery rate (FDR) control was applied (FDR ≤ 1%). Peptide-spectrum matches (PSMs) passing this threshold were filtered out, and a new MGF file was generated using the remaining unmatched spectra. In the third step, the unmatched spectra were searched against a customized small protein database (10 aa < length < 100 aa) with a more stringent FDR threshold of 0.1% to enhance the reliability of small protein identification. Key search parameters were as follows: protease was set to Trypsin, with full enzyme cleavage specificity; precursor mass tolerance was set to 10 ppm; the maximum number of modifications allowed per peptide was 3; and variable modifications included methionine oxidation.

### Prioritization of candidate AMPs for experimental validation

Potential AMPs were identified from the smORF dataset through a consensus screening strategy using dual-engine XAMP model. A candidate was advanced only if it received a prediction score >0.5 from both the XAMP-T and XAMP-E models. To ensure their clinical viability, additional selection criteria were implemented. ToxinPred^[33]^ and HemoPI^[34]^ were employed to exclude peptides exhibiting cytotoxicity or hemolytic activity, respectively. Subsequently, the modlAMP^[35]^ Python package was used to select peptides with a net charge exceeding +2. The AxPEP^[18]^ web server (https://app.cbbio.online/ampep/home) was then used to predict the minimum inhibitory concentrations (MICs) against Staphylococcus aureus and Escherichia coli. Peptides demonstrating both low predicted MIC values (≤ 20 μg/mL) and short lengths (≤ 40 aa) were prioritized for synthesis and validation, yielding seven final candidates.

### Materials

The AMPs used in this study were custom-synthesized by Sangon-Peptide Biotech (Ningbo) Co. Ltd., and their accurate molecular weights were determined by mass spectrometry. The purity of all peptides was determined by high-performance liquid chromatography (HPLC; column: kromasil C18, 5 μm, 4.6 mm × 150 mm; wavelength: 214 nm), and all purity was greater than 95%.

### MIC determination

The MICs of AMPs were determined using the broth microdilution method^[36]^ according to the Clinical and Laboratory Standards Institute (CLSI) 2025 guidelines^[37]^. The assay was validated by ensuring that the MIC of polymyxin B against the quality control strain *Escherichia coli* ATCC 25922 fell within the recommended CLSI range (0.25-2 μg/mL). The test organisms included Gram-negative bacteria *Escherichia coli* ATCC 8739, *Acinetobacter baumannii* ATCC 17978, *Pseudomonas aeruginosa* PAO1, *Klebsiella pneumoniae* ATCC 43816, and Gram-positive bacterium *Staphylococcus aureus* ATCC 6538. Briefly, two-fold serial dilutions of AMPs (256-0.5 μg/mL) were prepared in cation-adjusted Mueller-Hinton broth (CAMHB) 96-well plates. Each well, containing 50 µL of diluted AMP, was inoculated with 50 µL of a bacterial suspension prepared by adjusting cultures to 0.5 McFarland and diluting 1:100 in fresh CAMHB, yielding a final density of approximately 5×10^5^ CFU/mL in total volume of 100 µL. After incubation at 37 °C for 18-20 hours, the MIC was defined as the lowest peptide concentration that completely inhibited visible growth.

## Supporting information

Supplementary file 1.docx: Supplementary Table 1,2,4-6 and Supplementary Figure 1,2.

## Code availability

All code used in this study and the final trained models are provided in our public GitHub repository: https://github.com/Li-Lab-SJTU/XAMP.

## Supplementary information

The Supplementary information is available.

**Supplementary file 1.docx: Supplementary Table 1,2,4-6 and Supplementary Figure 1,2**.

Supplementary Table 1. Overview of trainset, testset and independent testset.

Supplementary Table 2. Model performance ablation study on different training datasets based on the Mix dataset (testset).

Supplementary Table 4. AMPs with spectral evidence in the metaproteome data.

Supplementary Table 5. The Information of AMPs used in this study.

Supplementary Figure 1. Taxonomic lineage of the bacterial-origin AMPs.

Supplementary Figure 2. Example peptide-spectrum matches (PSMs) for two AMPs identified in metaproteomic datasets.

**Supplementary file 2.xlsx: Supplementary Table 3**.

Supplementary Table 3. Deep-sea AMPs Database: 2,355 high-confidence peptides from consensus screening with dual-engine XAMP models.

## Acknowledgments

This work was supported by the Key Project for Computational Biology of Shanghai (grant no.: 23JS1400800), the National Natural Science Foundation of China (grant nos.: 32570783, 32170664, 42327805), the Fundamental Research Funds for the Central Universities (grant no.: YG2023ZD11), the Shanghai Pilot Program for Basic Research-Shanghai Jiao Tong University (grant no.: 21TQ1400201), and the National Key Research and Development Program of China (grant no.: 2023YFC2812800). The computations in this article were run on the Siyuan-1 cluster supported by the Center for High Performance Computing at Shanghai Jiao Tong University. The authors thank Zhuoqi Zheng and Jiayi Li of Shanghai Jiao Tong University for their valuable feedback on the manuscript.

## Authorship contributions

**Bairun Chen:** Investigation, Data Curation, Methodology, Statistical analysis, Writing - Original Draft, Visualization. **Xinyi Mou:** Investigation, Data Curation, Statistical analysis, Writing - Original Draft. **Zhuoxuan Song:** Investigation, Data Curation, Omics pipeline design and implementation. **Huaying Lin:** Methodology, Validation, Writing - Original Draft. **Tianyi Han:** Validation. **Runze Wang:** Resources. **Hong-Yu Ou:** Resources, Supervision. **Yu Zhang:** Supervision, Project Administration, Funding Acquisition. **Jing Li:** Conceptualization, Writing - Review & Editing, Supervision, Project Administration, Funding Acquisition. All authors have read and approved the final version of the manuscript.

