## Supplementary file 1.docx: Supplementary Table 1,2,4-6 and Supplementary Figure 1,2. for "XAMP: debiased dual-engine AI framework enables global discovery of antimicrobial peptides from deep-sea microbiomes"

**Supplementary Table 1. Overview of trainset, testset and independent testset.**

| Dataset | Positive | Negative | Total | *Remarks |
|---|---|---|---|---|
| Bacteria (trainset) | 526 | 5,950 | 6,476 | model suffix: "_bt" |
| Bacteria (testset) | 72 | 648 | 720 | |
| Unannotated (trainset) | 12,585 | 208,820 | 221,405 | model suffix: "_un" |
| Unannotated (testset) | 1,382 | 23,220 | 24,602 | |
| Mix (trainset) | 13,111 | 214,770 | 227,881 | model suffix: "_mix" |
| Mix (testset) | 1,454 | 23,868 | 25,322 | benchmark 1 |
| Xiao 1 (independent testset) | 757 | 2,386 | 3,143 | benchmark 2: removed the Xiao et al.'s benchmark dataset's AMPs existed in Unannotated (trainset) |
| Xiao 2 (independent testset) | 157 | 2,206 | 2,363 | benchmark 3: removed the Xiao et al.'s benchmark dataset's AMPs existed in Mix (trainset) |

**Supplementary Table 2. Model performance ablation study on different training datasets based on the Mix dataset (testset).**

| Model | Accuracy | Precision | AUC | AUPRC | FPR |
|---|---|---|---|---|---|
| XAMP_mix | 0.9795 | **0.9606** | **0.9715** | **0.8801** | **0.0017** |
| XAMP-T_mix | 0.9784 | 0.8807 | 0.9668 | 0.8472 | 0.0059 |
| XAMP-E_mix | **0.9822** | 0.9176 | 0.9706 | 0.8779 | 0.0041 |
| XAMP_un | 0.9798 | 0.9600 | 0.9616 | 0.8755 | 0.0017 |
| XAMP-T_un | 0.9794 | 0.8802 | 0.9584 | 0.8588 | 0.0062 |
| XAMP-E_un | 0.9808 | 0.8936 | 0.9675 | 0.8683 | 0.0055 |
| XAMP_bt | 0.9226 | 0.2900 | 0.7770 | 0.2272 | 0.0359 |
| XAMP-T_bt | 0.9052 | 0.2306 | 0.7420 | 0.1900 | 0.0566 |
| XAMP-E_bt | 0.9154 | 0.3341 | 0.8025 | 0.3745 | 0.0578 |
| iAMPCN | 0.2738 | 0.0701 | 0.7525 | 0.1945 | 0.7673 |
| c_AMPs | 0.9510 | 0.5610 | 0.8893 | 0.6786 | 0.0321 |
| Macrel | 0.9271 | 0.4097 | 0.8321 | 0.5494 | 0.0537 |
| APG | 0.7551 | 0.1722 | 0.8946 | 0.6248 | 0.2511 |

The optimal model for each evaluation metrics is given in boldface.

**Supplementary Table 4. AMPs with spectral evidence in the metaproteome data.**

| AMP sequence | Spectra count | Pride ID | Metagenomic evidence |
|---|---|---|---|
| PKMKTKKAAAKRFKITATGKLKHGVAGKRHRLMSHNSKYIRQNRGTKVASHADVARVVKKFLPYGL | 1 | PXD034421 | Yes |
| KRTFQPSNIKRKRNHGFRARMATKNGRKIVAARRAKGRKRLTA | 6 | PXD034421 | Yes |
| ARSLKKGPYIEHHLVKKVDVMNESGKKSVIKTWSRRSMISPDFVGHTFAVHNGNKFIPVFVTDNMVGHKLGEFAPTRNFRGHIAKKDKGKR | 3 | PXD034421 | No |
| ATKKAGGSSRNGRDSIGRRLGVKKFGGENVLAGNIIVRQRGTKFHPGNNVGIGKDHTIFATKNGKVAFKKTRIRTFISVIPA | 3 | PXD034421 | Yes |
| KRTFQPSVLKRKRTHGFRARMATANGRKVLARRRAKGRKVLSA | 2 | PXD034421 | Yes |

**Supplementary Table 5. The Information of AMPs used in this study.**

| ID | Length | Net Charge | Molecular Weight | Polypeptide Sequence (from N-terminus to C-terminus) |
|---|---|---|---|---|
| **1#_KR40** | 40 | +15 | 4829.69 | KRFGKYGRKVSWIVKRYGKYGRKLSSK VKGYGRYVRTVSW |
| **3#_RK23** | 23 | +11 | 2924.603 | RKFVKYFALYKLARKVLRRGRRV |
| **4#_IW38** | 38 | +15 | 4570.537 | IWKIRKKGKPNREKIGKIWKKIAKRYGK YGRTVSLNVK |
| **5#_KW39** | 39 | +13 | 4663.474 | KWYGKHGRKAGQIVIKYEKYGRKCWN VKGYGKYGRKTVK |
| **6#_GI39** | 39 | +16 | 4194.219 | GIKKGKKIPLSKLKAAAKKGGKLGKRA NLALTFRKMKKS |
| **7#_AR40** | 40 | +20 | 4806.837 | ARRKSWKKGKVFMKGRKKVRYIYPNG KKKGRKLVSASKRR |

**Supplementary Figure 1. Taxonomic lineage of the bacterial-origin AMPs. (A) Sankey diagram of taxonomic lineages of AMPs. G+, Gram-positive bacteria; G-, Gram-negative bacteria. (B) Bar plot of phylum-level distribution of AMPs.**

Bacterial-origin AMPs exhibited a taxonomic bias toward Gram-positive bacteria, with Gram-negative bacteria (e.g., Cyanobacteriota, Pseudomonadota) comprising only 14.1% of the dataset. At the phylum level, Bacillota was the most abundant source (312 AMPs), followed by Actinomycetota (48) and Pseudomonadota (26).

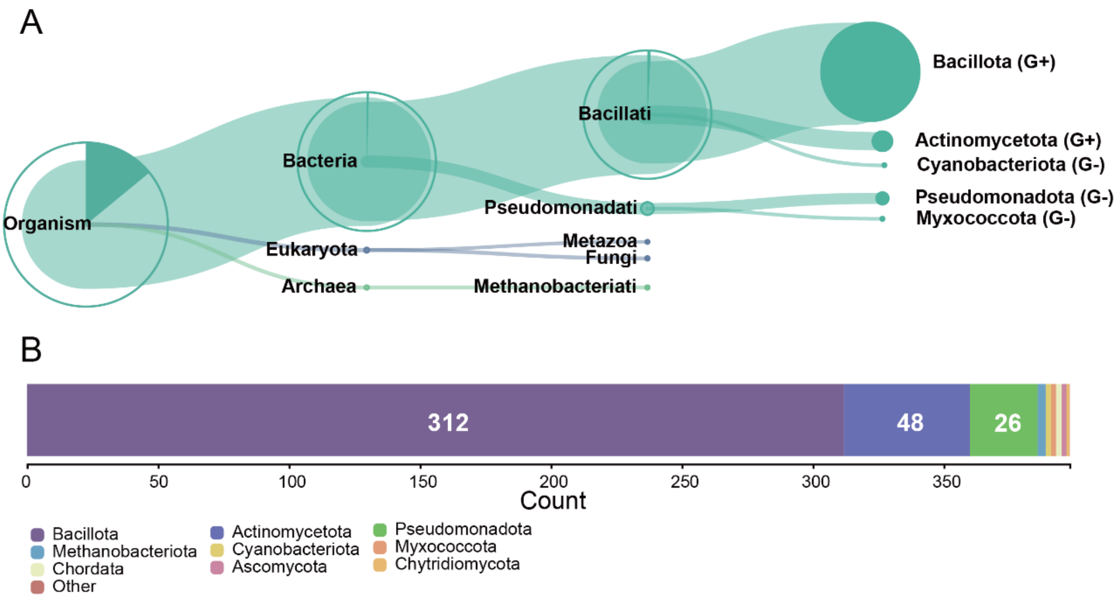

**Supplementary Figure 2. Example peptide-spectrum matches (PSMs) for two AMPs identified in metaproteomic datasets.**

**KRTFQPSVLKRKRTHGFRARMATANGRKVLARRRAKGRKVLSA**

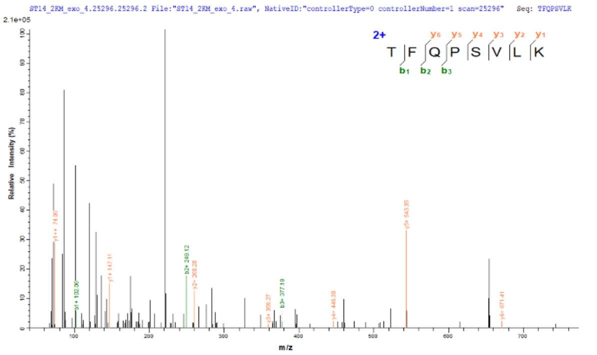

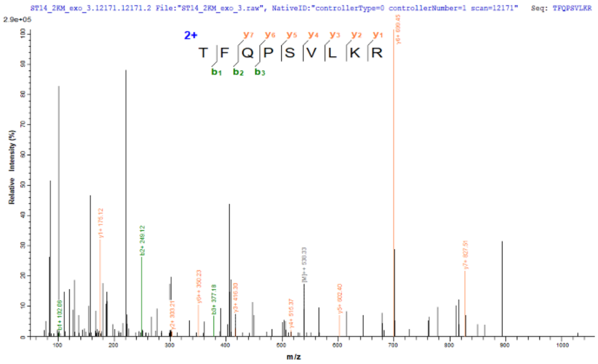

**PKMKTKKAAAKRFKITATGKLKHGVAGKRHRLMSHNSKYIRQNRGTKVASHADVARVVKKFLPYGL**

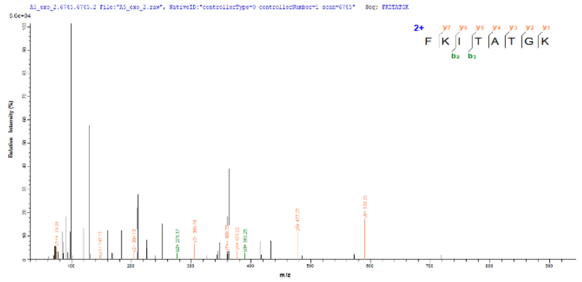
